# Genetic diversity and multiplicity of infection in *Fasciola gigantica* isolates of Pakistani livestock

**DOI:** 10.1101/789297

**Authors:** Zia Ur Rehman, Osama Zahid, Imran Rashid, Qasim Ali, Muhammad Haroon Akbar, Muhammad Oneeb, Wasim Shehzad, Kamran Ashraf, Neil D. Sargison, Umer Chaudhry

## Abstract

*Fasciola gigantica* liver flukes are responsible for over 3 billion US dollars of production loss annually in farmed livestock and cause widespread zoonotic disease. Nevertheless, the understating of the emergence and spread of the trematode species is poor. The multiplicity of *F. gigantica* infection and its spread is potentially influenced by multiple factors, including the abundance of suitable intermediate hosts, climatic conditions favoring the completion of the parasite’s lifecycle, and translocation of infected animals or free-living parasite stages between regions. Here we describe the development of a ‘tremabiome’ metabarcoding sequencing method to explore the numbers of *F. gigantica* genotypes per infection and patterns of parasite spread, based on genetic characteristics of the mitochondrial NADH dehydrogenase 1 (mt-ND-1) locus. We collected *F. gigantica* from three abattoirs in the Punjab and Balochistan provinces of Pakistan, and our results show a high level of genetic diversity in 20 *F. gigantica* populations derived from small and large ruminants consigned to slaughter in both provinces. This implies that *F. gigantica* can reproduce in its definitive hosts through meiosis involving cross- and self-breeding, as described in the closely related species, *Fasciola hepatica*. The genetic diversity between the 20 populations derived from different locations also illustrates the impact of animal movements on gene flow. Our results demonstrate the predominance of single haplotypes, consistent with a single introduction of *F. gigantica* infection in 85% of the hosts from which the parasite populations were derived. This is consistent with clonal reproduction in the intermediate snail hosts.

## 1. Introduction

Fascioliosis and fascioliasis are important neglected worldwide diseases of ruminant livestock and humans, respectively. *Fasciola* spp. not only cause billion of dollar production losses annually in farm animals (Aghayan et al., 2019; Mungube et al., 2006), but are also emerging food-borne zoonoses in humans according to the World Health Organization (WHO), with 2.4 million people infected and over 180 million at risk (Haseeb et al., 2002; Henok and Mekonnen, 2011). The genus *Fasciola* comprises of two important species, *Fasciola gigantica* and *Fasciola hepatica* (Mas-Coma et al., 2005). *F. gigantica* typically cause disease in tropical regions (Amor et al., 2011), whereas *F. hepatica* is present in temperate areas (Mazeri et al., 2017). Overlap between these two species has been seen in subtropical regions, along with the presence of intermediate forms (Ichikawa and Itagaki, 2010; Rokni et al., 2010).

Animals become infected with *Fasciola* spp. following the ingestion of herbage contaminated with infective metacercariae, and the parasites’ life cycles involve different snail species of *Galba truncatula* of Lymnaeidae family as intermediate hosts. Acquired parasite larvae penetrate the intestinal wall and migrate to the bile ducts of the definitive host, damaging the hepatic parenchyma and biliary tract (Geadkaew et al., 2011; Usip et al., 2014). Several potential factors may influence the emergence of *F. gigantica* infection. First, the seasonal presence of suitable intermediate hosts in areas with high rainfall, moderate temperatures, and humidity, and poor drainage (Claxton et al., 1997; Kaplan, 2001; Malek, 2018; Portugaliza et al., 2019; Rana et al., 2014; Rowcliffe and Ollerenshaw, 1960); resulting in high prevalences during certain months of the year (Phiri et al., 2005b; Rangel-Ruiz et al., 1999). Second, management stress, host immunity, and co-infection with intestinal nematodes; determining infection rates of *F. gigantica* (Ahmad et al., 2017; Elelu and Eisler, 2018; Phiri et al., 2005a). Third, the clonal expansion of *Fasciola* within its intermediate snail host; resulting in pasture contamination and subsequent host infection by metacercariae of the same genetic origin. This clonal expansion can also produce a genetic bottleneck effect when levels of infection in the snail populations are low (Beesley et al., 2017).

Over past few decades, high levels of animal movement have been reported in domestic ruminants in several European and Asian countries (Kelley et al., 2016; Vilas et al., 2012); hence genetic analyses are needed to understand the corresponding spread of *F. gigantica* infections and aid in the development of parasite control strategies (Hayashi et al., 2016). Animal movement patterns differ between farms and *F. gigantica* infects a wide range of hosts including farm and wild animals and humans. This potentially enables wildlife reservoirs of infection and cross-infection between different host species, complicating control strategies (Rojo-Vázquez et al., 2012).

Genetic relationships between parasite species and populations can be analysed using internal transcribed spacer region 2 (rDNA ITS-2) and mitochondrial nicotinamide adenine dinucleotide (NADH) dehydrogenase 1 (mt-ND-1) sequence data, respectively. Metabarcoding and deep amplicon sequencing methods and the Illumina Mi-Seq platform enable the study of large and diverse fluke populations, to show if infection has emerged recently in the host at a single point of time, or if burdens have been established repeatedly at different times, before spreading as a result of animal movement. Recently, we have used these methods to study the multiplicity of *Calicophoron daubneyi* infection in United Kingdom cattle herds. Our findings were consistent with multiple independent emergences of *C. daubneyi* infection; while the identification of common mt-COX-1 haplotypes in several populations across a range of geographic locations, highlighting the role of animal movements in the parasite’s spread (Sargison et al., 2019).

In this paper we describe a study using liver fluke parasites collected from cattle, buffalo, sheep and goats slaughtered in three geographically separated abattoirs in the Punjab and Balochistan provinces of Pakistan, with the following aims: i) to confirm the species identity of recovered *Fasciola* spp.; ii) to identify the presence of single or multiple genotypes per infection (multiplicity of infection) and iii) demonstrate the spread of *F. gigantica* mt-ND-1 haplotypes. The species identity of liver flukes was confirmed by deep amplicon sequencing of 483bp metabarcoding fragments of the rDNA ITS-2 region. Haplotype diversity in 20 *F. gigantica* populations, each derived from single infected ruminants consigned from different locations, was shown by deep amplicon sequencing of 311bp metabarcoding fragments of the mt-ND-1 locus. Split and network trees of the mt-ND-1 haplotypes were examined to show the multiplicity of *F. gigantica* infection and the patterns their spread, providing proof of concept for a novel approach to epidemiological studies of fasciolosis and fascioliasis, and validation of parasite control strategies.

## 2. Materials and methods

### 2.1. Sample collection and genomic DNA extraction

Three abattoirs were selected in the Punjab and Balochistan provinces of Pakistan, where there is a known high prevalence of fasciolosis. A total of 26 infected livers were collected from the three city abattoirs. Fourteen samples (buffalo=7, cattle=1, goat=6) were collected from Lahore, Punjab (31.5204° N, 74.3587° E), six samples (cattle=4, sheep=2) were collected from Murgha Kibzai, Balochistan (30.7384° N, 69.4136° E), and six samples (buffalo=3, sheep=3) were obtained from Loralai, Balochistan (30.3806° N, 68.5963° E). The livers were transported from abattoir to the laboratory on ice, where liver flukes were extracted from the biliary ducts by dissection. In total, 305 (183 from Punjab and 122 from Balochistan) individual flukes obtained from the populations. The flukes were identified as *Fasciola* spp. morphologically, thoroughly washed with phosphate-buffered saline (PBS), and preserved in 70% ethanol at −80 °C. For DNA extraction, a small tissue piece of around 2 mg was taken from the head of each worm, avoiding the possible contamination with any eggs. Each piece was rinsed twice for 5 min each, in a petri dish with distilled water (dH_2_O) and then lysed in 25 μl worm-lysis solution, prepared by mixing of 50 μl Proteinase K (10 mg/ml, New England BioLabs) and 50 μl 1M Dichloro Diphenyl Trichloroethane (DDT) in 1 ml of DirectPCR Lysis Reagent (Viagen). The lysates were then incubated at 60°C for 2 hrs, followed by 85°C for 15 min. Lysates were stored at −80°C until use. The abattoir-based study was approved by the Ethical Review Committee of the University of Veterinary and Animal Sciences Lahore Pakistan.

### 2.2. Metabarcoded sequencing approach

The overall scheme of the metabarcoding sequencing approach using Illumina MiSeq platform is shown in Fig. 1. The whole procedure was run in twice: first on the individual worms for species identification using the rDNA ITS-2 marker; then on pooled *F. gigantica* from each population, using the mt-ND-1 marker.

**Fig. 1.**
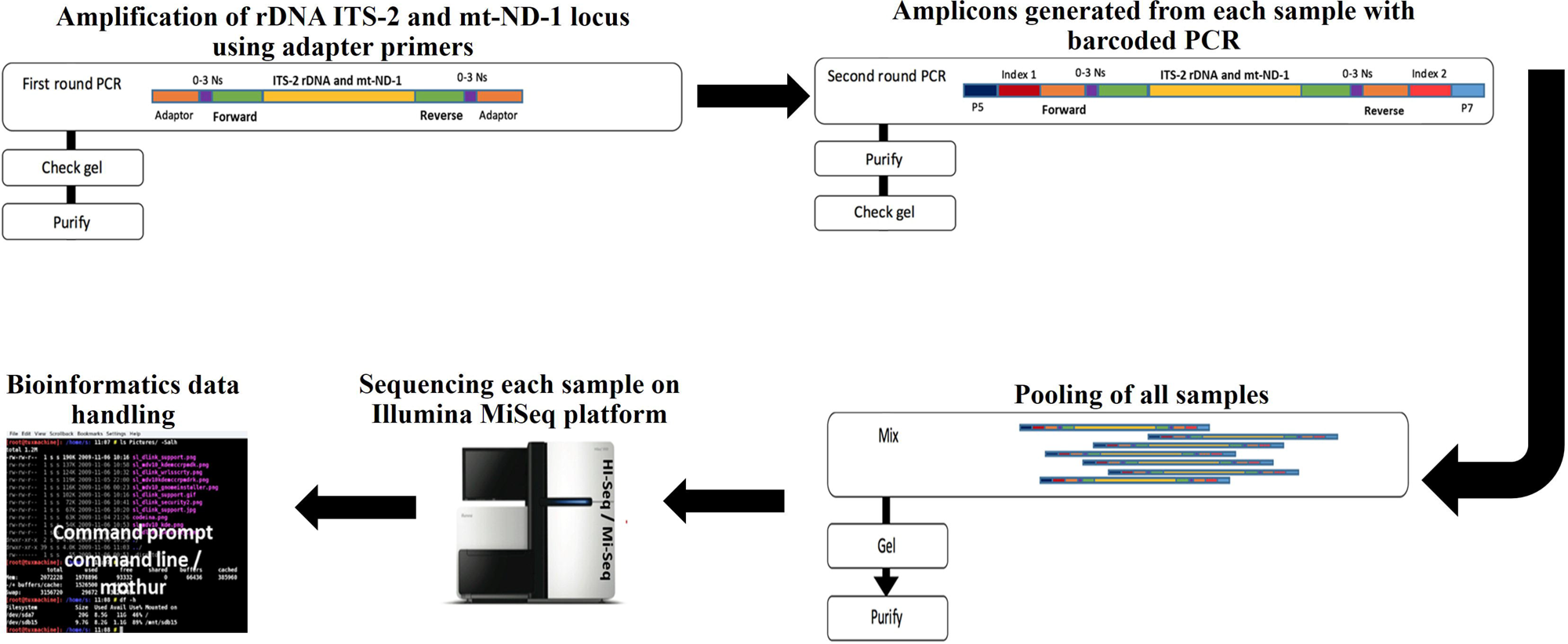
Schematic representation of the preparation and bioinformatics analysis of the metabarcoding sequencing libraries. In the first-round PCR amplification, overhanging primers were used to amplify the ITS-2 rDNA and mt-ND-1 fragments. The adapter base pairs provide the target sites for the primers used for sequencing the fragment. The random nucleotides (‘0-3N’s) are inserted between the adapter and the primers to offset the reading frame, thereby amplicons were sequenced to prevent the oversaturation of the MiSeq sequencing channels. The second-round PCR amplification was then performed using overhanging barcoded primers to bind to the adapter tags to add indices as well as the P5 and P7 regions required to bind to the MiSeq flow cell. The analysis of both rDNA ITS-2 and mt-ND-1 FASTQ files were performed in Mothur v1.39.5 software (Schloss et al., 2009) by using our modified Command Prompt pipeline and the standard operating procedures of Illumina MiSeq (Kozich et al., 2013).

#### 2.2.1. Adapter PCR amplification of rDNA ITS-2 and mt-ND-1 loci

The 1^st^ round PCR amplification was performed on 483bp fragments of the rDNA ITS-2 region, complementary to the 5.8s and 28s rDNA coding sequences, using universal sets of primers (Adlard et al., 1993; Chaudhry et al., 2016) and 311bp mt-ND-1 fragments, using newly developed primers (Supplementary Table S1, Fig. 1). The metabarcode adapters were added to the primers to allow their successive annealing, and the enhancement of the amplicons by adding up to three random nucleotides (N) between each primer and adapter, resulting in a total of four forward and four reverse primers (Supplementary Table S1). For both rDNA ITS-2 and mt-ND-1 loci, equal proportions of four forward and four reverse primers were mixed and used for the adapter PCR with the following conditions: 10 mM dNTPs, 10 μM forward and reverse adapter primers, 5X KAPA HiFi Fidelity buffer, 0.5 U KAPA HiFi Fidelity Polymerase (KAPA Biosystems, USA), 14 μl ddH_2_O and 1 μl of fluke lysate. The thermocycling conditions of the PCR reaction were as follows: 95°C for 2 min, followed by 35 cycles of 98°C for 20 sec, 65°C for 15 sec for ITS-2, and 60°C for 15 sec for ND-1, and 72°C for 15 sec, followed by a final extension at 72°C for 5 min. PCR products were then purified with AMPure XP Magnetic Beads (1X) according to the manufacturer’s instructions (Beckman Coulter).

#### 2.2.2. Barcoded PCR amplification of rDNA ITS-2 and mt-ND-1 loci

In the 2^nd^ round metabarcoded PCR, 16 forward (N501-N516) and 24 reverse (N701-N724) primers were used (Supplementary Table S2, Fig. 1) in a way that each sample contained a unique combination of forward and reverse primers. The barcoded PCR was performed with the following conditions: 10 mM dNTPs, 10 μM barcoded forward and reverse primers, 5X KAPA HiFi Fidelity buffer, 0.5 U KAPA HiFi Fidelity Polymerase (KAPA Biosystems, USA), 14μl ddH2O and 2μl of the first-round PCR product. The thermocycling conditions for this round of PCR were: 98°C for 45 sec, followed by 7 cycles of 98°C for 20 sec and 63°C for 20 sec, and final extension at 72°C for 2 min. The PCR products were purified with AMPure XP Magnetic Beads (1X) according to the manufacturer’s instructions (Beckman Coulter).

#### 2.2.3. Illumina Mi-Seq run, bioinformatics data handling and analysis

The bead-purified products from each sample were mixed to prepare a pooled library (Fig. 1) and measured with the KAPA qPCR library quantification kit (KAPA Biosystems, USA) before running it on the Illumina Mi-Seq Sequencer using a 500-cycle pair-end reagent kit (Mi-Seq Reagent Kits v2, MS-103-2003) at a concentration of 15nM with addition of 15% Phix Control v3 (Illumina, FC-11-2003). During the post-run processing, Mi-Seq splits all sequences by samples using the barcoded indices to produce FASTQ files (Supplementary Table S2). The analysis of both rDNA ITS-2 and mt-ND-1 FASTQ files were performed in Mothur v1.39.5 software (Schloss et al., 2009), using our modified Command Prompt pipeline (Sargison et al., 2019)., and the standard operating procedures of Illumina Mi-Seq (Kozich et al., 2013).

For both rDNA ITS-2 and mt-ND-1, the raw paired-end reads were analysed to combine the two sets of reads for each parasite population using make.contigs command, which requires ‘stability.files’ as an input. The ‘make.contigs’ command extracts sequence and quality score data from FASTQ files, creating complements of the reverse and forward reads and joins them into contigs. It simply aligns the pairs of sequence reads and compares the alignments to identify any positions where the two reads disagree. Next, there was a need to remove any sequences with ambiguous bases using the ‘screen.seqs’ command. The alignment of the above dataset was performed with a *F. gigantica* and *F. hepatica* rDNA ITS-2 and *F. gigantica* mt-ND-1 reference sequence taxonomy library created from the NCBI database (for more details Supplementary Data S1; Section 2.4) based on where the sequences start and end corresponding with the primer set (Supplementary Table S1). The ‘align.seqs’ command was adapted for the reference sequence taxonomy library to be aligned with rDNA ITS-2 and mt-ND-1 Illumina Mi-Seq dataset. To confirm that these filtered sequences overlap the same region of the reference sequence taxonomy library, the ‘screen.seqs’ command was run to show the sequences ending at the 483bp rDNA ITS-2 and 311bp mt-ND-1 position.

At this stage, the rDNA ITS-2 analysis was completed by classifying the sequences into either of the two groups (*F. gigantica* or *F. hepatica*), using the classify.seqs command and creating the taxonomy file by using the summary.tax command. Overall, thousands of rDNA ITS-2 reads were generated from the data set of 305 individual worms from 26 populations. The presence of each species was calculated by dividing the number of sequences reads of each worm by the total number of reads.

For mt-ND-1 analysis, once all sequences were classified as *F. gigantica* mt-ND-1 based on the ‘screen.seqs’ command, a count list of the consensus sequences of each population was created using the unique.seqs command, followed by the use of pre.cluster command to look for sequences having up to two differences and to merge them in groups based on their abundance. Chimeras were identified and removed using the chimera.vsearch commands. The count list was then used to create the FASTQ files of the consensus sequences of each population using the split.groups command (for more details Supplementary Data S2). The consensus sequences of *F. gigantica* mt-ND-1 were analysed separately in Geneious v9.0.1 software (Biomatters Ltd, New Zealand) (Kearse et al., 2012) using the MUSCLE alignment tool to remove the polymorphisms occurring once only as being artifacts due to sequencing errors as previously described (Sargison et al., 2019). These aligned consensus sequences were then imported into the FaBox 1.5 online tool to collapse the sequences that showed 100% base pair similarity after corrections into a single haplotype. The frequency of all the haplotypes present in the total and at each population level was calculated by dividing the number of sequences reads of each population by the total number of reads.

### 2.3. Network and Split tree analysis

A network tree was produced based on a neighbor-joining algorithm using Network 4.6.1 software (Fluxus Technology Ltd), built on a sparse network with the epsilon parameter is set to zero default. A split tree was created in the SplitTrees4 software (Huson and Bryant, 2005) by using the UPGMA method in the Jukes-Cantor model of substitution. The appropriate model of nucleotide substitutions for UPGMA analysis was selected by using the jModeltest 12.2.0 program (Posada, 2008). A maximum-likelihood tree of was also constructed in Mega 6.0 (Tamura et al., 2013) based on a neighbor-joining algorithm using HKY+G+I model. The branch supports were obtained by 1000 bootstraps of the data.

Genetic diversity was calculated for all the generated sequences within and between populations by using the DnaSP 5.10 software package (Librado and Rozas, 2009), and the following values were obtained: Haplotype diversity (Hd), the number of segregating sites (S), nucleotide diversity (π), the mean number of pairwise differences (k), the mutation parameter based on an infinite site equilibrium model, and the number of segregating sites (θS).

## 3. Results

### 3.1. Consensus sequence taxonomy library preparation of F. gigantica and F. hepatica rDNA ITS-2 and mt-ND-1 loci

A total of 27 NCBI GeneBank sequences of the *Fasciola* rDNA ITS-2 locus (*F. gigantica*=*14*, *F. hepatica*=*13*) and 167 NCBI GeneBank sequences of the *Fasciola* mt-ND-1 locus (*F. gigantica*=*67*, *F. hepatica*=*100*) were identified by BLAST search (for more details Supplementary Data S1). The sequences were analysed separately in Geneious v9.0.1 software (Biomatters Ltd, New Zealand) (Kearse et al., 2012) using the MUSCLE alignment tool. The construction of the maximum-likelihood tree demonstrate distinct clustering of rDNA ITS-2 and mt-ND-1 loci of *F. gigantica* and *F. hepatica* (Fig. 2A & B).

**Fig. 2.**
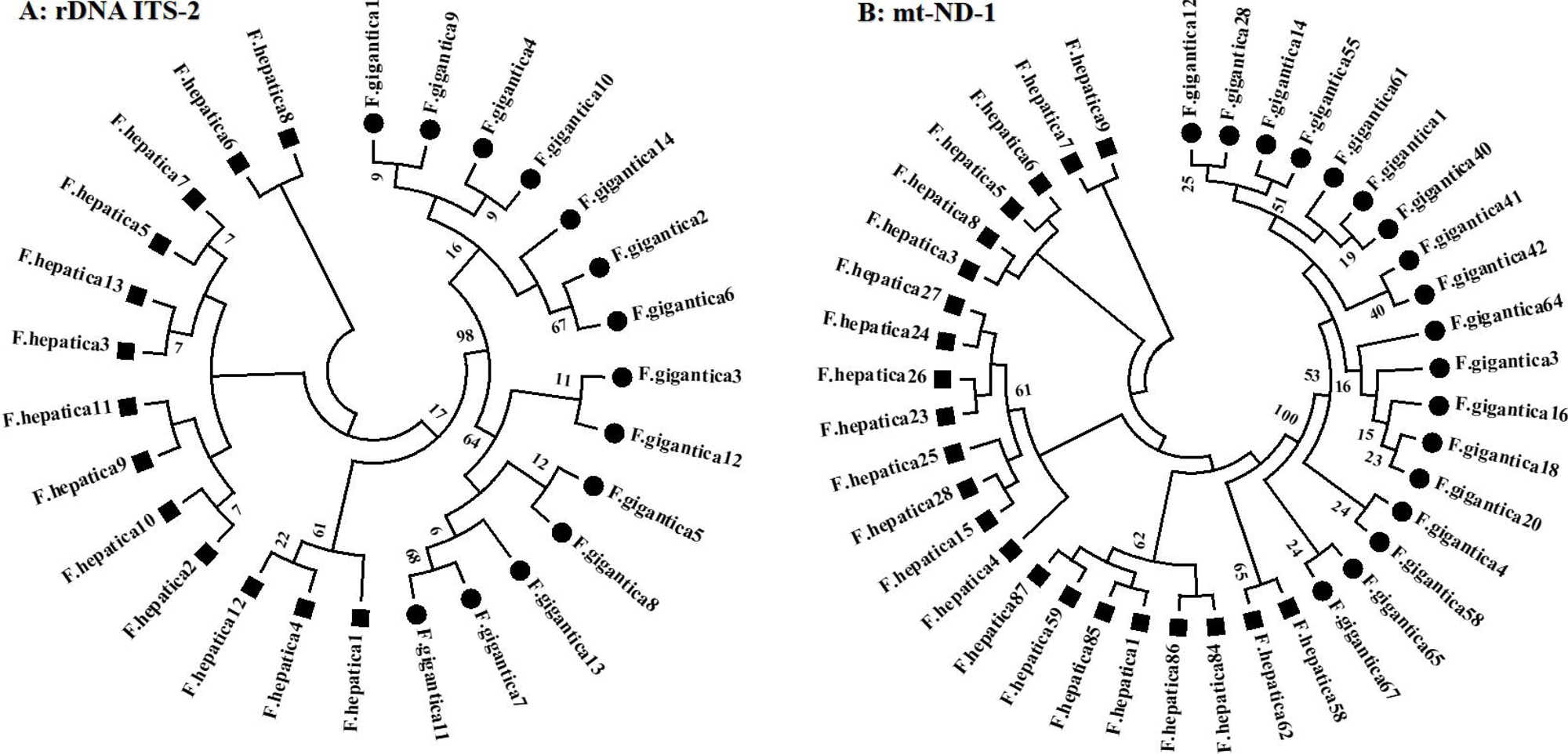
The maximum-likelihood tree was obtained from the BLAST searched *F. gigantica* and *F. hepatica* rDNA ITS-2 and mt-ND-1 loci. The NCBI GeneBank sequences were first aligned on the MUSCLE tool of the Geneious v9.0.1 software. (A) Twenty-seven sequences of the rDNA ITS-2 were identified among *F. gigantica and F. hepatica* species. (B) Forty unique sequences of the mt-ND-1 were identified among hundred and sixty-seven *F. gigantica and F. hepatica* species. The neighbor-joining algorithm (Kimura 2 and HKY+G+I-parameter model) was computed with 1000 bootstrap replicates using MEGA5 software created by Biomatters. Each species was identified with different shades bars.

### 3.2. Confirmation of species identity and co-infection by rDNA ITS-2 sequence analysis

A total 305 individual worms, comprising of 26 populations, were run through the Illumina MiSeq platform, out of which 238 (77.6%) were confirmed to be *F. gigantica*, 56 (18.7%) were *F. hepatica* and 11 (3.7%) showed heterozygous sequence reads (Table 1). All 183 worms collected in Punjab from buffalo, cattle, and goats were *F. gigantica*. In contrast, 122 worms collected from Balochistan were both *F. gigantica* (42.2%) and *F. hepatica* (48.3%), as well as the intermediate form (9.5%); except the buffalo populations contained only *F. gigantica* (Table 1).

**Table 1.** Species identification of individual worms. from twenty-six Fasciola populations were performed. Each population represents the flukes collected from a single host. The table is divided into two sections based on the province from where the samples were collected.

| Populations | Host | Area | Province | Total flukes | *F. gigantica* | *F. hepatica* | Heterozygous |
| --- | --- | --- | --- | --- | --- | --- | --- |
| P2B | Buffalo | Lahore | Punjab | 16 | 16 | - | - |
| P4B | Buffalo | Lahore | Punjab | 15 | 15 | - | - |
| P7B | Buffalo | Lahore | Punjab | 8 | 8 | - | - |
| P13B | Buffalo | Lahore | Punjab | 20 | 20 | - | - |
| P14B | Buffalo | Lahore | Punjab | 20 | 20 | - | - |
| P15B | Buffalo | Lahore | Punjab | 20 | 20 | - | - |
| P16B | Buffalo | Lahore | Punjab | 20 | 20 | - | - |
| P26C | Cattle | Lahore | Punjab | 20 | 20 | - | - |
| P1G | Goat | Lahore | Punjab | 4 | 4 | - | - |
| P6G | Goat | Lahore | Punjab | 16 | 16 | - | - |
| P9G | Goat | Lahore | Punjab | 7 | 7 | - | - |
| P10G | Goat | Lahore | Punjab | 6 | 6 | - | - |
| P11G | Goat | Lahore | Punjab | 4 | 4 | - | - |
| P12G | Goat | Lahore | Punjab | 7 | 7 | - | - |
| P3B | Buffalo | Loralai | Balochistan | 11 | 11 | - | - |
| P5B | Buffalo | Loralai | Balochistan | 12 | 12 | - | - |
| P8B | Buffalo | Loralai | Balochistan | 7 | 7 | - | - |
| P17C | Cattle | Murgha Kipzai | Balochistan | 12 | 3 | 6 | 3 |
| P18C | Cattle | Murgha Kipzai | Balochistan | 12 | 8 | 2 | 2 |
| P19C | Cattle | Murgha Kipzai | Balochistan | 18 | 8 | 7 | 3 |
| P20C | Cattle | Murgha Kipzai | Balochistan | 20 | 3 | 16 | 1 |
| P21S | Sheep | Murgha Kipzai | Balochistan | 15 | 3 | 11 | 1 |
| P22S | Sheep | Murgha Kipzai | Balochistan | 3 | - | 3 | - |
| P23S | Sheep | Loralai | Balochistan | 1 | - | 1 | - |
| P24S | Sheep | Loralai | Balochistan | 3 | - | 3 | - |
| P25S | Sheep | Loralai | Balochistan | 8 | - | 7 | 1 |
| **Total** |  |  |  | **305** | **238** | **56** | **11** |

### 3.3. Haplotype distribution of F. gigantica mt-ND-1 locus

The haplotype frequencies present at the individual population level were analysed separately from 20 selected *F. gigantica* populations (Fig. 3). Only three populations (P8B, P12G, and P18C) each containing between 7 and 8 worms, showed relatively high frequencies of multiple haplotypes. These three populations contained 7, 3, and 13 haplotypes (Fig. 3). In contrast, single haplotypes predominated in 17 populations, each containing between 4 and 20 worms (Fig. 3). Three of these populations (P13B, P7B, and P17C) contained 6 haplotypes; six populations (P2B, P14B, P15B, P16B, P19B, and P26C) had 5 haplotypes; three populations (P4B, P6G, and P10G) had 4 haplotypes; four populations (P1G, P9G, P3B, and P11G) had 3 haplotypes; and one population (P5B) contained only one haplotype (Fig. 3).

**Fig. 3.**
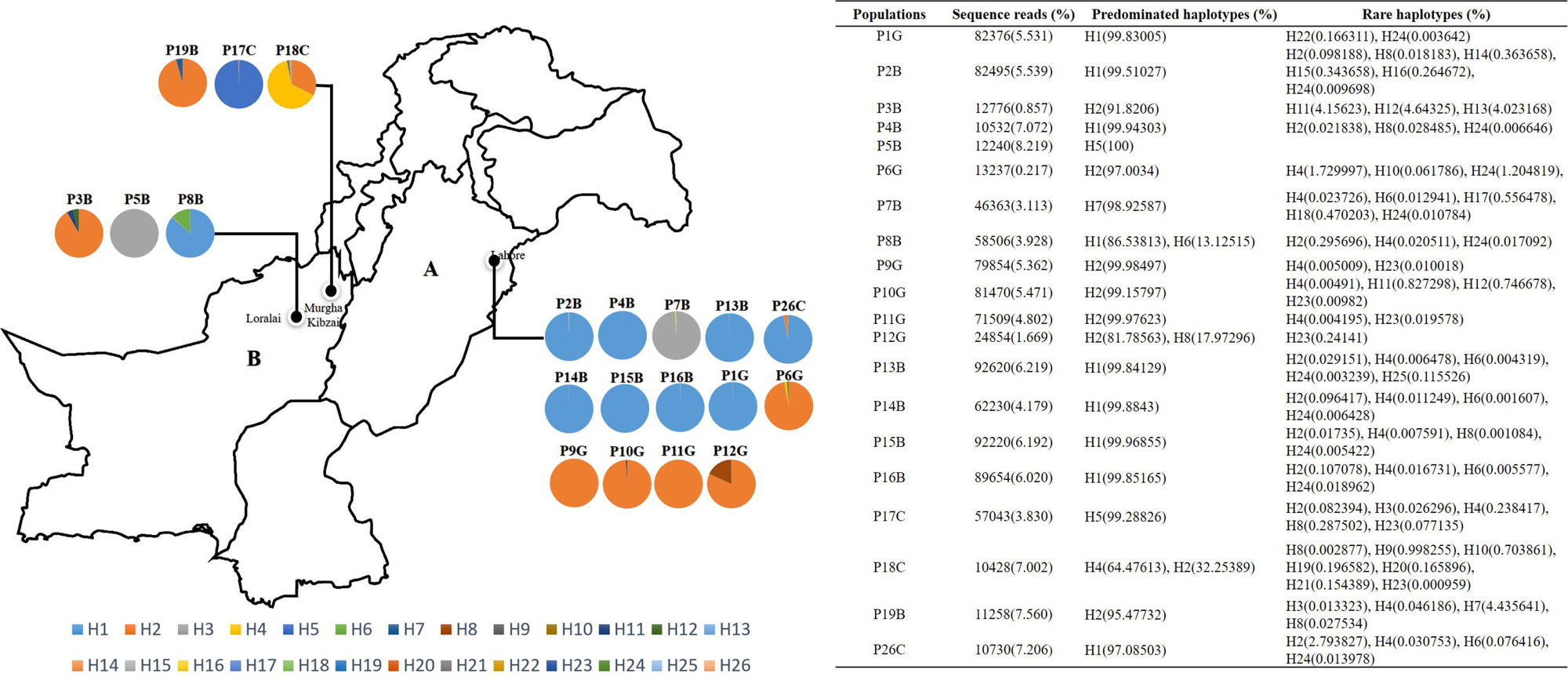
Relative allele frequencies of twenty *F. gigantica* populations from Punjab (A) and Balochistan (B) provinces of Pakistan. Each pie chart displays the individual population, and each haplotype in the pie chart is represented by a different color codes. The pie chart circle represents the population distribution and their frequency comes from each of the twenty-six haplotypes as indicated on the insert table. The populations are also linked to the location from where they were collected.

Twenty-six haplotypes of the *F. gigantica* mt-ND-1 locus were identified among all 20 populations. The split tree analysis shows at least two distinct clades (Fig. 4). Six haplotypes (H13, H26, H4, H19, H9, and H20) in Clade I, were present only in Balochistan. The remaining 20 haplotypes were in clade II, with 8 haplotypes (H17, H21, H14, H10, H12, H5, H7, and H6) predominating in Balochistan. Contrastingly, the six haplotypes (H17, H21, H14, H10, H12, H5, H7, and H6) were more common in Punjab. In addition, four equally shared haplotypes (H2, H3, H11, and H23) were present in clade II.

**Fig. 4.**
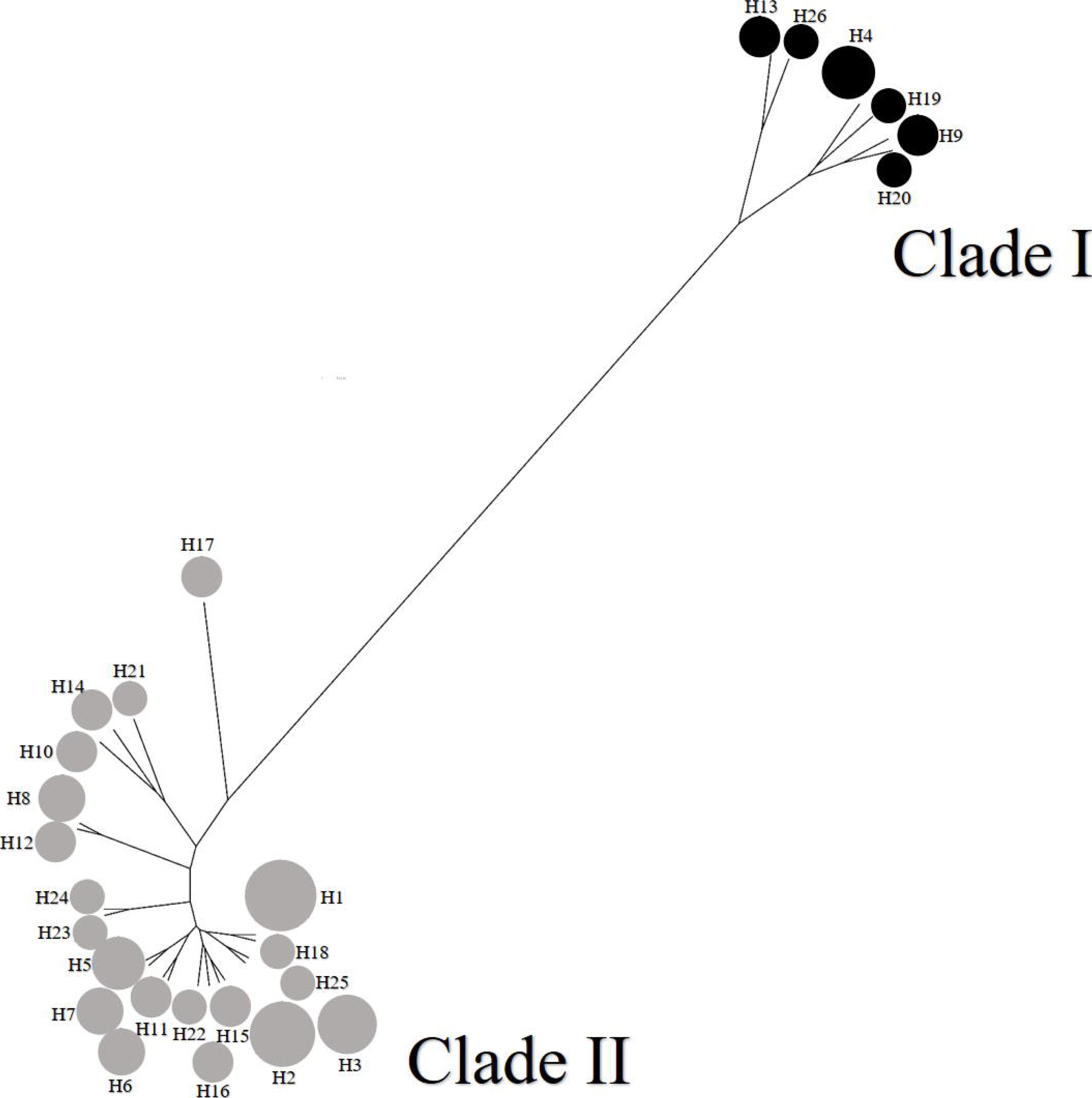
Split tree of twenty-six mt-ND-1 haplotypes generated from twenty *F. gigantica* populations. The tree was constructed with the UPGMA method in the Jukes-Cantor model of substitution in the SplitsTrees4 software. The appropriate model of nucleotide substitutions for UPGMA analysis was selected by using the jModeltest 12.2.0 program. The pie chart circles in the tree represent the haplotypes in two different clades and the size of each circle is proportional to the frequency in the population.

### 3.4. Phylogenetic analysis of F. gigantica mt-ND-1 locus

The network and maximum likelihood trees were produced to examine the phylogenetic relationship between 26 different haplotypes of the *F. gigantica* mt-ND-1 locus (Fig. 5, Supplementary Fig. S1). Sixteen haplotypes were comparatively rare as they collectively made less than 5% of the total reads. Five of these haplotypes (H15, H16, H18, H22, and H25) were only found in Punjab, and they were present in four populations; whereas 11 haplotypes (H5, H7, H9, H12, H13, H14, H17, H19, H20, H21, and H26) were present in Balochistan, originating from 5 populations (Fig. 5, Supplementary Fig. S1). The remaining 10 haplotypes were shared between Punjab and Balochistan, making more than 95% of the total reads. Three of these haplotypes (H1, H8, and H24) predominated in Punjab and were present in 13 populations in total, including 9 populations from Punjab and 4 from Balochistan (Fig. 5, Supplementary Fig. S1). Three haplotypes (H4, H6, and H10) predominated in Balochistan and were present in 14 populations in total, including 5 from Balochistan and 9 from Punjab (Fig. 5, Supplementary Fig. S1). In contrast, four haplotypes (H2, H3, H11, and H23) were equally shared between Punjab and Balochistan (Fig. 5, Supplementary Fig. S1); these were present in a total of 19 populations, 13 coming from Punjab and 6 from Balochistan.

**Fig. 5.**
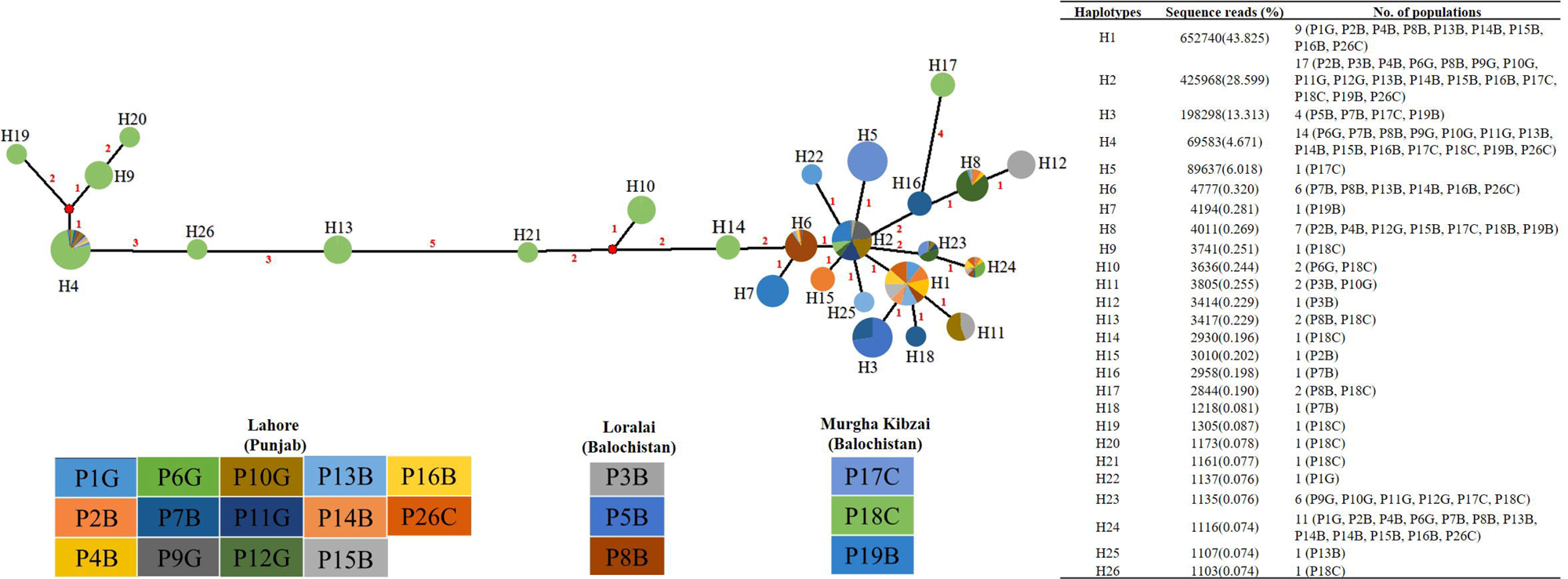
Network tree of twenty-six mt-ND-1 haplotypes sequenced from twenty *F. gigantica* populations. The tree was generated with the Neighbour Joining method in the Network 4.6.1 software (Fluxus Technology Ltd.). All unnecessary median vectors and links were removed with the star contractions. Each pie chart displays the individual haplotype, and each population in the pie chart is represented by a different color codes. The pie chart circle represent the haplotype distribution and their frequency comes from each of the twenty populations as indicated on the insert table. The color of each haplotype circle represents the percentage of sequence reads generated per population shown in the insert table. The number of mutations (in red) separating adjacent sequence nodes was indicated along with the connecting branches and the length of the lines connecting the haplotypes is proportional to the number of nucleotide changes.

### 3.5. Genetic diversity of F. gigantica mt-ND-1 locus

The data show a high level of genetic diversity at both haplotype and nucleotide levels (Table 2), with the values of haplotype diversity ranging from 0.727 to 0.960 and nucleotide diversity from 0.00643 to 0.03118 within different populations. The overall values between populations were 0.917 and 0.02466 for haplotype and nucleotide diversity, respectively. This analysis confirms that Pakistani *F. gigantica* populations have a high level of genetic diversity.

**Table 2:**
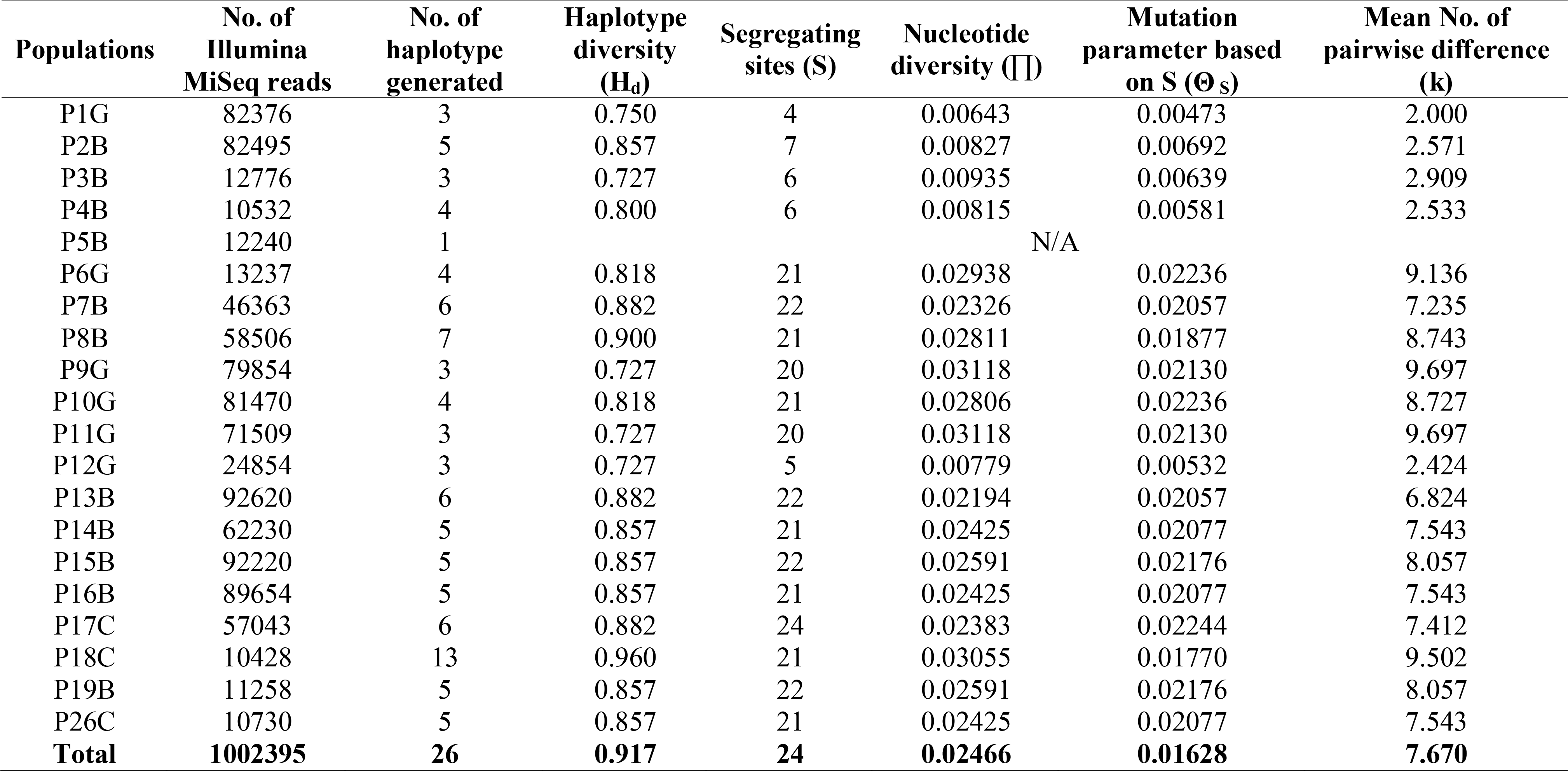
Genetic diversity estimation of mt-ND-1 haplotypes identified from twenty populations of Fasciola gigantica in Pakistan. The table shows these values both with populations and the total between populations.

## 4. Discussion

Next-generation genomic resources have potential applications to investigate parasite species dynamics, coinfections, hybridisation, multiplicity of infection and the level of gene flow, as well as evaluation of population responses to drug treatments (Sargison et al., 2019). Various PCR methods [(RFLP)-PCR, quantitative (q)PCR, multiplex PCR, SSCP-PCR, Sanger sequencing] have been described to amplify the mt-ND-1 and rDNA ITS-2 regions for sequence determination (Ai et al., 2011), but these are low throughput, hence relatively expensive, and potentially error-prone. In contrast, high throughput deep amplicon sequencing or metabarcoded DNA derived from parasite populations using the Illumina MiSeq platform is relatively low-cost and potentially less error-prone. The method has transformed the study of haemoprotozoan parasites (Chaudhry et al., 2019) and nematode parasites (Avramenko et al., 2015); and has the potential to open new areas of research in the study of trematode parasite communities. We have used this method to study the multiplicity of *C. daubneyi* infection in the United Kingdom (Sargison et al., 2019). The use of primers binding to conserved sites and analysis of up to 600 bp sequence reads allows trematode species (the ‘tremabiome’) to be determined. The use of barcoded primers allows a large number of samples to be pooled and sequenced in a single Mi-Seq run, making the technology suitable for high-throughput analysis. By multiplexing the barcoded primer combinations, it is possible to run 384 samples at once on a single Illumina Mi-Seq flow cell, helping to reduce the cost (Chaudhry et al., 2019; Sargison et al., 2019). In the present study, using high throughput deep amplicon sequencing of metabarcoded DNA, we first confirm the presence of *F. gigantica* and then identify the prevalence of single or multiple haplotypes per infection (multiplicity of infection) and demonstrating the spread of *F. gigantica* alleles.

*Fasciola* spp. are cosmopolitan parasites that are widely distributed throughout the world (Mas-Coma, 2004). The prevalence of *F. gigantica* is highest in tropical and subtropical regions while *F. hepatica* is more common in temperate areas where sheep and cattle are raised (Mas-Coma et al., 2014). Pakistan has an agriculture-based economy with livestock being an integral part. There are several reports of *Fasciola* infections in Pakistan, mostly implying that *F. hepatica* is most commonly identified in small and large ruminants (Ashraf et al., 2014; Ijaz et al., 2009; Shahzad et al., 2012). A few reports indicate the presence of both *F. hepatica* and *F. gigantica*, with intermediate forms found (Akhtar et al., 2012; Iqbal et al., 2007). However, all of these reports are based on egg and adult morphology with limited molecular confirmation of species identity. In the present study of the dynamics of *Fasciola* spp. in Pakistan, ITS-2 rDNA sequence data confirmed the presence of *F. hepatica*, *F. gigantica,* and intermediate forms, with *F. gigantica* being the predominant species (Table 1), concurring with a previous study (Chaudhry et al., 2016).

The predominance of single mt-ND-1 haplotypes in 17 of 20 (85%) *F. gigantica* populations suggests a single genetic emergence of infection in the host from which the parasites were derived. In contrast, the presence of high proportions of multiple mt-ND-1 haplotypes in just 3 of 20 (15%) populations implies multiple emergences (Fig. 3). The clonal lineage of the haplotypes in most of the *F. gigantica* populations is similar to the situation that has been described in *F. hepatica* (Beesley et al., 2017; Vilas et al., 2012), implying similar importance of asexual reproduction of the intermediate snail hosts to the population genetics of the two species. The clonal lineage might arise because the life cycle of *F. gigantica* involves uneven disease transmission intensity and, or, abundance of intermediate hosts; as has been described for *F. hepatica,* where single miracidia infect the *G. truncatula* mud snails, giving rise to multiple, genetically identical cercariae (Beesley et al., 2017). Then, after the release of clonal lineage cercariae by the intermediate hosts, there may be an aggregation of infective metacercaria, with little mixing on the pasture, prior to the ingestion by the definitive host. These conditions would be more likely if the prevalence of infection in the intermediate hosts is low, and snail habitats are small and isolated, as is likely the case in Pakistan, where long-term muddy regions tend to be found in association with irrigation pumps or ponds. An alternative explanation for the clonal lineage of the *F. gigantica* could be a little population bottlenecking arising due to the influences of the wide range of climatic conditions throughout the year on intermediate snail host infection (Rana et al., 2014).

Our results confirm a high level of genetic diversity in *F. gigantica* (Table 2), implying the capacity for reproduction in the definitive host through meiosis during cross-breeding. A similar situation has been described in *F. hepatica* infection (Beesley et al., 2017; Vilas et al., 2012; Zintl et al., 2015). However, a low level of infection (Dar et al., 2011; Rondelaud et al., 2011) and clonal lineage in snails could give rise to a potential low population bottlenecking effect, raising questions about how genetic diversity might be maintained in *F. gigantica*. For example, could snails be infected with multiple miracidia and subsequently shed cercariae with different alleles, or could diversity be maintained by long survival of metacercariae on the pasture?.

The movement of livestock is frequent in Pakistan and contributes to the high levels of parasitic gene flow (Ali et al., 2018). In the present study, ten haplotypes were spread between populations. Three of those haplotypes predominated in the Punjab and three in the Balochistan provinces. In contrast, four haplotypes were equally shared between the populations from both provinces. This suggests that animal movement plays a key role in maintaining a high level of gene flow in *F. gigantica*, for example translocating animals to a new region could introduce new parasites as well as exposing the definitive hosts to different residents parasite populations.

In summary, we used a mt-ND-1 marker to provide first insights into the genetic diversity and multiplicity of *F. gigantica* infection in Pakistan. Our findings suggest that most of the hosts were predominantly infected with parasites of identical mt-ND-1 haplotypes, consistent with clonal multiplication within the snails. A high level of genetic diversity was seen in *F. gigantica* isolated from naturally infected small and large ruminants, showing that sexual reproduction with cross-fertilisation occurs within the definitive host. The most common mt-ND-1 haplotypes were identified in several populations across a range of geographic locations, highlighting the role of animal movements in the spread of infectious disease. Our study provides proof of concept for a method that could be used to investigate host-parasite relationships and to examine influences of intermediate hosts, co-infections and climate change on the epidemiology of *F. gigantica*; with reference to the development of sustainable parasite control strategies.

## Acknowledgment

The study was financially supported by Roslin Institute using facilities funded by Biotechnology and Biological Sciences Research Council (BBSRC). Work at the University of Veterinary and Animal Science Pakistan uses facilities funded by the Higher Education Commission of Pakistan. The authors of this study would like to thank Dr. Talat Naseer Pasha-Vice-Chancellor of the University of Veterinary and Animal Science Lahore Pakistan for his great support in the arrangements for sample collections.

## Supplementary Figure Legends

**Supplementary Fig. S1.** Maximum likelihood tree of twenty-six mt-ND-1 haplotypes sequenced from twenty *F. gigantica* populations from two provinces of Pakistan as indicated on the insert table. The tree was constructed in Mega 6.0 (Tamura et al., 2013) based on a neighbor-joining algorithm using HKY+G+I model. The tree was rooted with the corresponding mt-ND-1 sequence of *F. hepatica* (4P01770.1), and the branch supports were obtained by 1000 bootstraps of the data. The colored rectangles and circles indicate the populations in which each haplotype was identified as indicated by the color key in the insert table.

