## Supplementary figures and images for "Genetic diversity and multiplicity of infection in *Fasciola gigantica* isolates of Pakistani livestock"

### Supplementary Figure S1

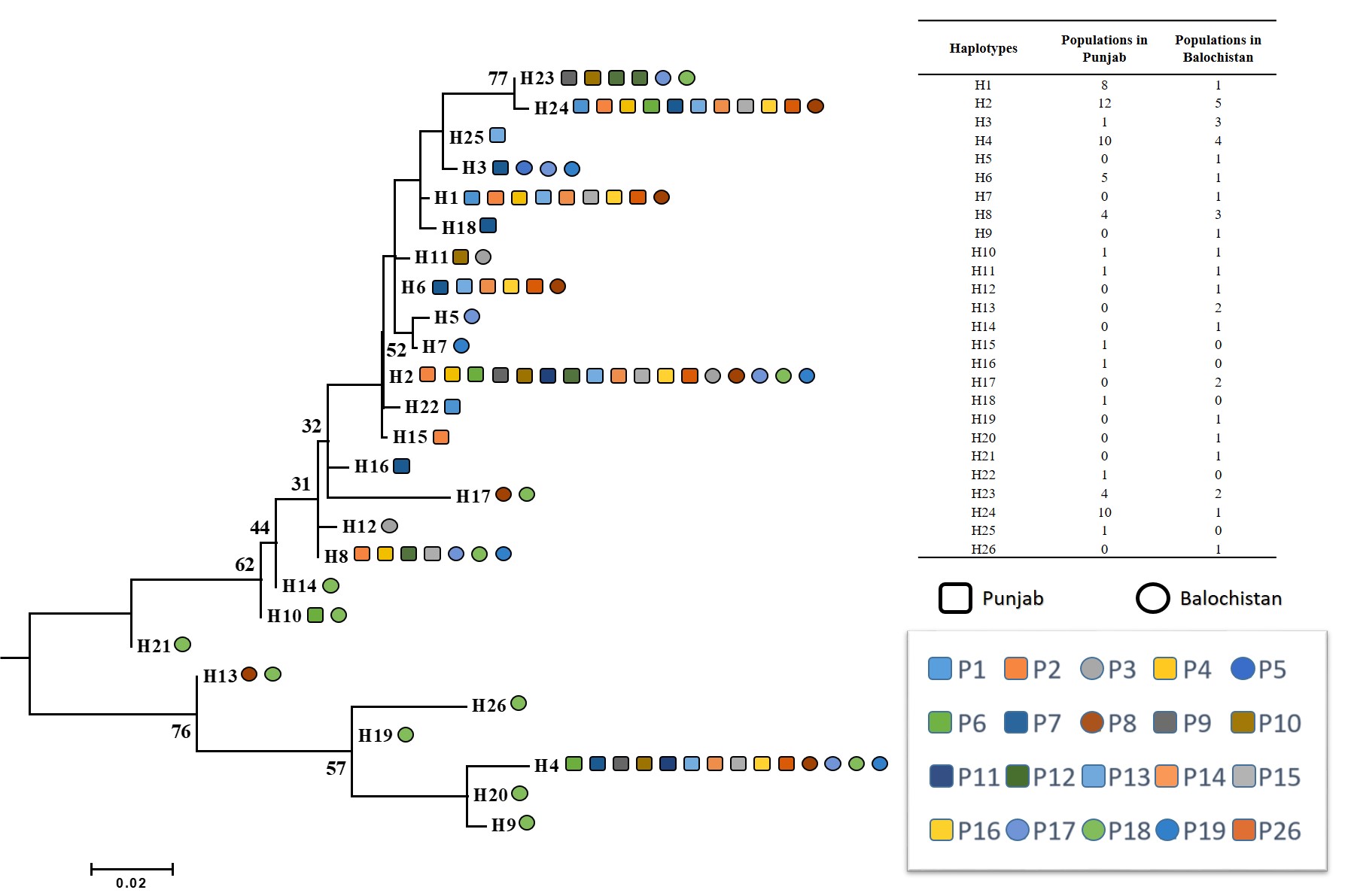
