## Supplementary Table S1 for "Genetic diversity and multiplicity of infection in *Fasciola gigantica* isolates of Pakistani livestock"

**Supplementary Table S1**. ITS-2 and mt-ND-1 primers. Primer sequences for the amplification of Fasciola rDNA ITS-2 and mt-ND-1. Forward and reverse primer sets are underlined, N’s are bolded, and adapters are in italic format.

| **Sequences (5'-3')** | **Primer name** | **Target region** | **Direction** |
| --- | --- | --- | --- |
| *TCGTCGGCAGCGTCAGATGTGTATAAGAGACAG*GGTGGATCACTCGGCTCG*T*G | AD_For | rDNA ITS-2 | Forward |
| *TCGTCGGCAGCGTCAGATGTGTATAAGAGACAG***N**GGTGGATCACTCGGCTCG*T*G | AD_For-1N | rDNA ITS-2 | Forward |
| *TCGTCGGCAGCGTCAGATGTGTATAAGAGACAG***NN**GGTGGATCACTCGGCTCG*T*G | AD_For-2N | rDNA ITS-2 | Forward |
| *TCGTCGGCAGCGTCAGATGTGTATAAGAGACAG***NNN**GGTGGATCACTCGGCTCG*T*G | AD_For-3N | rDNA ITS-2 | Forward |
| *GTCTCGTGGGCTCGGAGATGTGTATAAGAGACAG*TTCCTCCGCTTAGTGATAT*G*C | AD_Rev | rDNA ITS-2 | Reverse |
| *GTCTCGTGGGCTCGGAGATGTGTATAAGAGACAG***N**TTCCTCCGCTTAGTGATAT*G*C | AD_Rev-1N | rDNA ITS-2 | Reverse |
| *GTCTCGTGGGCTCGGAGATGTGTATAAGAGACAG***NN**TTCCTCCGCTTAGTGATAT*G*C | AD_ Rev-2N | rDNA ITS-2 | Reverse |
| *GTCTCGTGGGCTCGGAGATGTGTATAAGAGACAG***NNN**TTCCTCCGCTTAGTGATAT*G*C | AD_ Rev-3N | rDNA ITS-2 | Reverse |
| *TCGTCGGCAGCGTCAGATGTGTATAAGAGACAG*GTTTAAGTTTGTGTTTTT*T*C | UN2_For | mt-ND-1 | Forward |
| *TCGTCGGCAGCGTCAGATGTGTATAAGAGACAG***N**GTTTAAGTTTGTGTTTTT*T*C | UN2_For-1N | mt-ND-1 | Forward |
| *TCGTCGGCAGCGTCAGATGTGTATAAGAGACAG***NN**GTTTAAGTTTGTGTTTTT*T*C | UN2_For-2N | mt-ND-1 | Forward |
| *TCGTCGGCAGCGTCAGATGTGTATAAGAGACAG***NNN**GTTTAAGTTTGTGTTTTT*T*C | UN2_For-3N | mt-ND-1 | Forward |
| *GTCTCGTGGGCTCGGAGATGTGTATAAGAGACAG*CACCATAACTCCCCCAAAC*C*A | ND1_Rev | mt-ND-1 | Reverse |
| *GTCTCGTGGGCTCGGAGATGTGTATAAGAGACAG***N**CACCATAACTCCCCCAAAC*C*A | ND1_Rev-1N | mt-ND-1 | Reverse |
| *GTCTCGTGGGCTCGGAGATGTGTATAAGAGACAG***NN**CACCATAACTCCCCCAAAC*C*A | ND1_Rev-2N | mt-ND-1 | Reverse |
| *GTCTCGTGGGCTCGGAGATGTGTATAAGAGACAG***NNN**CACCATAACTCCCCCAAAC*C*A | ND1_Rev-3N | mt-ND-1 | Reverse |
