## Supplementary Table S1 for "Genetic diversity and multiplicity of infection in *Fasciola gigantica* isolates of Pakistani livestock"

**Supplementary Table S2.** List of barcoded primers. Sequences for forward and reverse barcoded primers (Nextera XT Index Kit v2). Index sequences are highlighted in bold. The primers were used in a way that each sample had a unique forward-reverse combination. Sequences came from Oligonucleotide sequences © 2018 Illumina, Inc. All rights reserved.

| Primer | Sequence, 5’-3’ |
| --- | --- |
| S501 | AATGATACGGCGACCACCGAGATCTACAC**TAGATCGC**TCGTCGGCAGCGTC |
| S502 | AATGATACGGCGACCACCGAGATCTACAC**CTCTCTAT**TCGTCGGCAGCGTC |
| S503 | AATGATACGGCGACCACCGAGATCTACAC**TATCCTCT**TCGTCGGCAGCGTC |
| S504 | AATGATACGGCGACCACCGAGATCTACAC**AGAGTAGA**TCGTCGGCAGCGTC |
| S505 | AATGATACGGCGACCACCGAGATCTACAC**GTAAGGAG**TCGTCGGCAGCGTC |
| S506 | AATGATACGGCGACCACCGAGATCTACAC**ACTGCATA**TCGTCGGCAGCGTC |
| S507 | AATGATACGGCGACCACCGAGATCTACAC**AAGGACTA**TCGTCGGCAGCGTC |
| S508 | AATGATACGGCGACCACCGAGATCTACAC**CTAAGCCT**TCGTCGGCAGCGTC |
| S510 | AATGATACGGCGACCACCGAGATCTACAC**CGTCTAAT**TCGTCGGCAGCGTC |
| S511 | AATGATACGGCGACCACCGAGATCTACAC**TCTCTCCG**TCGTCGGCAGCGTC |
| S512 | AATGATACGGCGACCACCGAGATCTACAC**TCGACTAG**TCGTCGGCAGCGTC |
| S513 | AATGATACGGCGACCACCGAGATCTACAC**TTCTAGCT**TCGTCGGCAGCGTC |
| S514 | AATGATACGGCGACCACCGAGATCTACAC**CCTAGAGT**TCGTCGGCAGCGTC |
| S515 | AATGATACGGCGACCACCGAGATCTACAC**GCGTAAGA**TCGTCGGCAGCGTC |
| S516 | AATGATACGGCGACCACCGAGATCTACAC**CTATTAAG**TCGTCGGCAGCGTC |
| N701 | CAAGCAGAAGACGGCATACGAGAT**TAAGGCGAG**TCTCGTGGGCTCGG |
| N702 | CAAGCAGAAGACGGCATACGAGAT**CGTACTAGG**TCTCGTGGGCTCGG |
| N703 | CAAGCAGAAGACGGCATACGAGAT**AGGCAGAAG**TCTCGTGGGCTCGG |
| N704 | CAAGCAGAAGACGGCATACGAGAT**TCCTGAGCG**TCTCGTGGGCTCGG |
| N705 | CAAGCAGAAGACGGCATACGAGAT**GGACTCCTG**TCTCGTGGGCTCGG |
| N706 | CAAGCAGAAGACGGCATACGAGAT**TAGGCATGG**TCTCGTGGGCTCGG |
| N707 | CAAGCAGAAGACGGCATACGAGAT**GTGTGTAGG**TCTCGTGGGCTCGG |
| N708 | CAAGCAGAAGACGGCATACGAGAT**CAGAGAGGG**TCTCGTGGGCTCGG |
| N709 | CAAGCAGAAGACGGCATACGAGAT**GCTAGGGTG**TCTCGTGGGCTCGG |
| N710 | CAAGCAGAAGACGGCATACGAGAT**CGAGGCTGG**TCTCGTGGGCTCGG |
| N711 | CAAGCAGAAGACGGCATACGAGAT**AAGAGGCAG**TCTCGTGGGCTCGG |
| N712 | CAAGCAGAAGACGGCATACGAGAT**GTAGAGGAG**TCTCGTGGGCTCGG |
| N713 | CAAGCAGAAGACGGCATACGAGAT**GCTCATGAG**TCTCGTGGGCTCGG |
| N714 | CAAGCAGAAGACGGCATACGAGAT**ATCTCAGGG**TCTCGTGGGCTCGG |
| N715 | CAAGCAGAAGACGGCATACGAGAT**ACTCGCTAG**TCTCGTGGGCTCGG |
| N716 | CAAGCAGAAGACGGCATACGAGAT**GGAGCTACG**TCTCGTGGGCTCGG |
| N717 | CAAGCAGAAGACGGCATACGAGAT**GCGTAGTAG**TCTCGTGGGCTCGG |
| N718 | CAAGCAGAAGACGGCATACGAGAT**CGGAGCCTG**TCTCGTGGGCTCGG |
| N719 | CAAGCAGAAGACGGCATACGAGAT**TACGCTGCG**TCTCGTGGGCTCGG |
| N720 | CAAGCAGAAGACGGCATACGAGAT**ATGCGCAGG**TCTCGTGGGCTCGG |
| N721 | CAAGCAGAAGACGGCATACGAGAT**TAGCGCTCG**TCTCGTGGGCTCGG |
| N722 | CAAGCAGAAGACGGCATACGAGAT**ACTGAGCGG**TCTCGTGGGCTCGG |
| N723 | CAAGCAGAAGACGGCATACGAGAT**CCTAAGACG**TCTCGTGGGCTCGG |
| N724 | CAAGCAGAAGACGGCATACGAGAT**CGATCAGTG**TCTCGTGGGCTCGG |
